# Genes lost during the transition from land to water in cetaceans highlight genomic changes involved in aquatic adaptations

**DOI:** 10.1101/521617

**Authors:** Matthias Huelsmann, Nikolai Hecker, Mark S. Springer, John Gatesy, Virag Sharma, Michael Hiller

**Affiliations:** Max Planck Institute of Molecular Cell Biology and Genetics, 01307 Dresden, Germany; Max Planck Institute for the Physics of Complex Systems, 01187 Dresden, Germany; Center for Systems Biology Dresden, 01307 Dresden, Germany; Department of Evolution, Ecology, and Organismal Biology, University of California, Riverside, CA 92521, USA; Division of Vertebrate Zoology and Sackler Institute for Comparative Genomics, American Museum of Natural History, New York, NY 10024, USA

**Keywords:** gene loss, cetaceans, aquatic adaptations

## Abstract

The transition from land to water in whales and dolphins (cetaceans) was accompanied by remarkable anatomical, physiological and behavioral adaptations. To better understand the genomic changes that occurred during this transition, we systematically screened for protein-coding genes that were inactivated in the ancestral cetacean lineage. We discovered genes whose loss is likely beneficial for cetaceans by reducing the risk of thrombus formation during diving (*F12*, *KLKB1*), improving the fidelity of oxidative DNA damage repair (*POLM*), and protecting from oxidative stress-induced lung inflammation (*MAP3K19*). Additional gene losses may reflect other diving-related adaptations, such as enhanced vasoconstriction during the diving response (mediated by *SLC6A18*) and altered pulmonary surfactant composition (*SEC14L3*), while loss of *SLC4A9* relates to a reduced need for saliva in aquatic environments. Finally, the complete loss of melatonin synthesis and receptor genes (*AANAT*, *ASMT*, *MTNR1A*/*B*) may have been a precondition for the evolution of unihemispheric sleep. Our findings suggest that some genes lost in the ancestral cetacean lineage may have been involved in adapting to a fully-aquatic lifestyle.

## Introduction

The ancestors of modern cetaceans (whales, dolphins and porpoises) transitioned from a terrestrial to a fully-aquatic lifestyle during the Eocene about 50 Mya (*1*). This process constitutes one of the most dramatic macroevolutionary transitions in mammalian history and was accompanied by profound anatomical, physiological and behavioral transformations that allowed cetaceans to adapt and thrive in the novel habitat (*2*). Remarkable changes in cetacean anatomy include streamlined bodies and loss of body hair to reduce drag during swimming, a much thicker skin that lacks sweat and sebaceous glands and has enhanced physical barrier properties, a thick layer of blubber for insulation, the loss of hindlimbs after propulsion by the tail flukes evolved, and reduced olfactory and gustatory systems, which became less important in water (*3*). To efficiently store and conserve oxygen for prolonged breath-hold diving, cetaceans developed a variety of adaptations. These adaptations include increased oxygen stores that result from large blood volumes and elevated concentrations of hemoglobin, myoglobin and neuroglobin in blood, muscle and brain tissue, respectively, a high performance respiratory system that allows rapid turnover of gases at the surface, and a flexible ribcage that allows the lung to collapse at high ambient pressure (*4*–*7*).

Comparative analysis of cetacean genomes has provided important insights into the genomic determinants of cetacean traits and aquatic specializations (*8*). Several studies revealed patterns of positive selection in genes with roles in the nervous system, osmoregulation, oxygen transport, blood circulation, protein and lipid digestion, or bone microstructure (*9*–*15*). An adaptive increase in myoglobin surface charge likely permitted a high concentration of this oxygen transport and storage protein in cetacean muscles (*16*). In addition to patterns of positive selection, the loss (inactivation) of protein-coding genes is associated with derived cetacean traits. For example, cetaceans have lost a large number of olfactory and vomeronasal receptors, olfactory signal transduction genes, taste receptors, and hair keratin and epidermal genes (*17*–*27*). Furthermore, all or individual cetacean lineages lost the ketone body synthesizing enzyme *HMGCS2* (*28*), the sebaceous gland-expressed *MC5R* gene (*29*), the non-shivering thermogenesis gene *UCP1* (*30*), the protease *KLK8* that plays distinct roles in the skin and hippocampus (*31*), short-and long-wave sensitive opsin genes (*32, 33*), and the antiviral genes *MX1* and *MX2* (*34*). During evolution, gene loss can be a consequence of relaxed selection on a function that became obsolete, but also can be a mechanism for adaptation (*35*). For example, the loss of the erythrocyte-expressed *AMPD3* gene in the sperm whale, one of the longest and deepest diving cetacean species, is likely beneficial by enhancing oxygen transport (*36*). The loss of the elastin-degrading protease *MMP12* may have contributed to ‘explosive exhalation’, and the loss of several epidermal genes (*GSDMA*, *DSG4*, *DSC1*, *TGM5*) likely contributed to hair loss and the remodeling of the cetacean epidermal morphology (*36*).

Due to an extensive series of intermediate fossils, the shift from a terrestrial to a fully-aquatic environment is one of the best characterized macroevolutionary transitions in mammalian evolution (*8*). However, the important genomic changes that occurred during this transformation remain incompletely understood. Since recent work has shown that the loss of ancestral protein-coding genes is an important evolutionary force, we conducted a systematic screen for genes that were inactivated on the stem Cetacea branch, i.e., after the split between Cetacea and Hippopotamidae, but before the split between Odontoceti (toothed whales) and Mysticeti (baleen whales). This revealed a number of gene losses that are associated with the evolution of adaptations to a fully-aquatic environment.

## Results and Discussion

### Screen for coding genes that were inactivated in the cetacean stem lineage

To investigate the contribution of gene inactivation to the evolution of adaptations to a fully-aquatic environment, we systematically searched for protein-coding genes exhibiting inactivating mutations that are shared between the two extant cetacean clades, odontocetes and mysticetes (exemplified in Figure 1A). The most parsimonious hypothesis for such shared inactivating mutations is that they occurred before the split of these two clades in the common ancestral branch of Cetacea. In our screen, we also included genes that are partially or completely deleted in both odontocetes and mysticetes and exhibit shared deletion breakpoints, indicating a single loss in cetacean stem lineage. In our genomic screen, odontocetes were represented by the bottlenose dolphin, killer whale and sperm whale (*37*–*39*), and mysticetes were represented by the common minke whale (*11*). We excluded genes that are lost in more than three terrestrial mammal species as these genes are likely not involved in the adoption of a fully-aquatic lifestyle. To precisely identify genes that were inactivated during the transition from land to water in the cetacean stem lineage, we made use of the recently-sequenced genome of the common hippopotamus (*40*), a semiaquatic mammal that along with the pygmy hippopotamus are the closest living relatives to cetaceans (*41*), and considered only genes with no detected inactivating mutations in the hippopotamus. To detect gene-inactivating mutations, we employed a comparative approach that makes use of genome alignments to search for mutations that disrupt the protein’s reading frame (stop codon mutations, frameshifting insertions or deletions, deletions of entire exons) and mutations that disrupt splice sites (*36*). We excluded olfactory receptor and keratin-associated genes since losses in these large gene families have previously been reported (*18*, *19*, *24*, *25*). This resulted in a set of 85 genes that exhibit shared inactivating mutations in odontocetes and mysticetes, most of which have not been reported before (Table S1).

**Figure 1:**
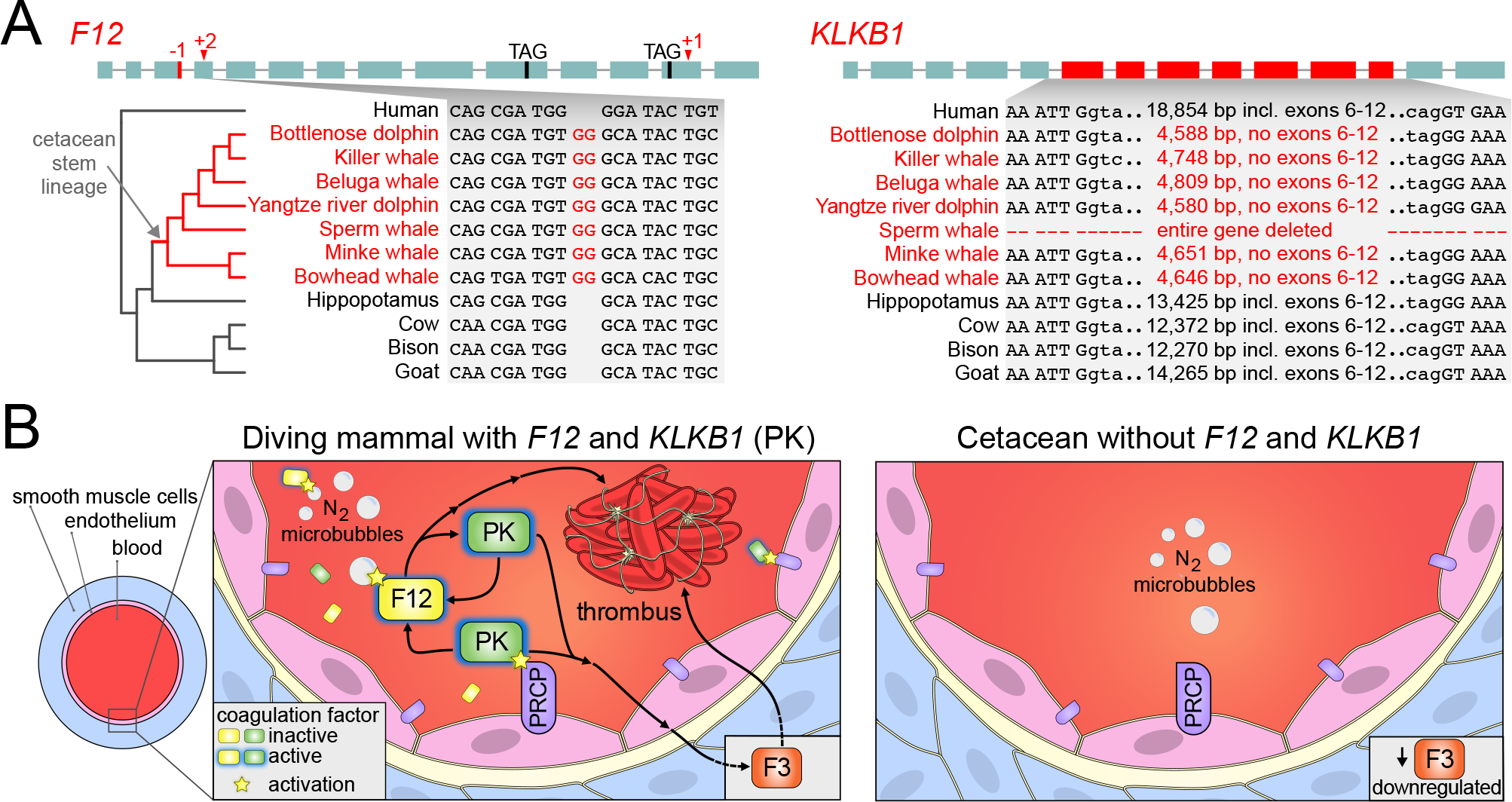
Key coagulation factors that promote thrombosis were lost in the cetacean stem lineage. (A)*F12* and *KLKB1* were lost in the cetacean stem lineage, consistent with previous findings (*11*, *39*, *43*, *45*). Boxes illustrate coding exons, superimposed with those gene-inactivating mutations that are shared among odontocetes and mysticetes, and thus likely occurred before the split of these lineages. The inset shows one representative inactivating mutation. Shared breakpoints imply that the deletion of *KLKB1* coding exons 6-12 occurred in the cetacean stem lineage (intronic bases adjacent to exons 5 and 13 are in lower case letters). All inactivating mutations in both genes are shown in Figures S2 and S3. (B) Left: *F12* encodes a zymogen that auto-activates by contact with a variety of surfaces, which likely include nitrogen microbubbles that form during breath-hold diving (*47*, *54*, *55*). *KLKB1* encodes another zymogen that can be activated to plasma kallikrein (PK) by either activated F12 or by the endothelial membrane-associated endopeptidase PRCP (*46*). PK in turn can activate F12. Both activated F12 and PK proteases promote thrombosis formation (*46*). Right: Gene knockouts in mice suggest that loss of *F12* and *KLKB1* has no major effect on wound sealing, but protects from thrombus formation via different mechanisms (*49*, *52*, *56*). While loss of *KLKB1* protects from thrombosis by reducing the expression of coagulation factor III (F3) (*57*), loss of *F12* prevents activation on nitrogen microbubbles during diving. Since a vasoconstriction-induced reduction in blood vessel diameters and nitrogen microbubble formation increase the risk of thrombosis for frequent divers, the loss of both genes was likely beneficial for cetaceans.

For these 85 genes, we performed additional analyses to confirm evolutionary loss in the cetacean stem lineage. First, inactivating mutations shared between the four cetaceans used in the genomic screen imply that other species that descended from their common ancestor should share these mutations. We tested this by manually inspecting the genomes of two additional odontocetes (Yangtze river dolphin (*23*) and beluga whale (*42*)) and an additional mysticete (bowhead whale (*43*)), which confirmed the presence of shared inactivating mutations. It should be noted that the presence of the same inactivating mutations in independent genome assemblies of multiple related species excludes the possibility that these mutations are sequencing or assembly errors, which is corroborated by DNA sequencing reads (Figure S1). Second, we manually inspected the genome alignments, which revealed no evidence for an undetected functional copy of these genes in cetaceans. Taken together, these analyses show that these genes were inactivated on the stem Cetacea branch, i.e. after the split between Cetacea and Hippopotamidae, but before the split between Odontoceti and Mysticeti.

We intersected the 85 genes with functional annotations of their human and mouse orthologs (Table S1) and performed a literature search. This revealed a number of genes that are likely related to aquatic adaptations (*F12*, *KLKB1*, *POLM*, *MAP3K19*, *SEC14L3*, *SLC6A18*, *SLC4A9*, *AANAT*) by being involved in thrombosis, repair of oxidative DNA damage, oxidative stress-induced lung inflammation, renal amino acid transport, saliva secretion, and melatonin synthesis. For these genes, we also verified that they not only have an intact reading frame in the common hippopotamus, but also in the pygmy hippopotamus, the only other extant species in the family Hippopotamidae. In the following, we describe how the loss of these genes likely relates to adaptations to a fully-aquatic environment.

### Loss of coagulation-associated factors and reduced thrombus formation

Diving results in a systemic response, consisting of a decrease in heart rate (bradycardia) and reduced peripheral blood flow, which is achieved by contraction of endothelial smooth muscle cells (peripheral vasoconstriction) (*44*). A frequent vasoconstriction-induced reduction in blood vessel diameter during diving increases the risk of thrombus (blood clot) formation. Our screen detected the loss of two blood coagulation-associated factors, *F12* (coagulation factor XII) and *KLKB1* (kallikrein B1). Several shared inactivating mutations show that both genes were lost in the cetacean stem lineage (Figures 1A, S2, S3). While the loss of these genes in various cetacean species was noted before (*11*, *39*, *43*, *45*), the mechanisms by which these two gene losses likely protect from thrombus formation during diving have not been described.

*F12* initiates thrombus formation via the contact activation system (CAS) (*46*). *F12* encodes a zymogen that auto-activates upon encountering a variety of foreign or biological surfaces (*47*). The activated zymogen functions as a serine protease that engages in a reciprocal activation cycle with the serine protease encoded by *KLKB1*, resulting in platelet activation and the formation of a blood clot (*46*, *48*). Consistently, knockout or knockdown of *F12* protects various mammals from induced thrombosis, but importantly did not impair wound sealing after blood vessel injury (hemostasis) (*49*–*52*). Eliminating CAS-based coagulation by inactivating *F12* may have been especially advantageous for cetaceans, as nitrogen microbubbles, which readily form in the blood upon repeated breath-hold diving (*53*), may act as foreign F12-activating surfaces entailing harmful thrombus formation (*54*, *55*) (Figure 1B).

The *KLKB1*-encoded zymogen prekallikrein is activated by proteolytic cleavage to form the serine protease plasma kallikrein (PK). Similar to the knockout of *F12*, the knockout of *KLKB1* in mice also granted protection from induced thrombosis while only slightly prolonging wound sealing (*52*, *56*). Interestingly, thrombosis protection in *KLKB1* knockout mice is mediated by a CAS-independent mechanism (*57*). *KLKB1* knockout leads to reduced levels of bradykinin, the main target of PK, which in turn leads to reduced expression of coagulation factor III (also called tissue factor, *F3*) (*57*), a key initiator of the blood coagulation cascade. Reduction of coagulation factor III alone is sufficient to reduce the risk of thrombosis (*58*). In addition to being activated by F12, prekallikrein can be activated by the endothelial membrane-associated endopeptidase PRCP (*59*). Evidently, this type of activation should happen more frequently in a diving cetacean, where constricted blood vessels increase the proximity of prekallikrein and PRCP (Figure 1B). Moreover, the activity of PRCP is pH dependent and peaks in slightly acidified plasma (*59*), a condition found in diving cetaceans (*60*).

In summary, all cetaceans are deficient in two key factors that promote thrombosis, but largely do not affect wound sealing. The risk of thrombus-induced occlusion of blood vessels is higher for frequent divers, as smaller blood vessel diameters and nitrogen microbubble formation during diving both increase the likelihood of F12 or prekallikrein activation (Figure 1B). Since inactivating *F12* or *KLKB1* reduces the risk of thrombus formation via different and likely additive mechanisms, both gene losses were potentially advantageous for stem cetaceans. Consistent with this, previous studies found that several genes involved in blood clotting or platelet formation evolved under positive selection in cetaceans (*38*, *61*).

### Loss of a DNA repair gene and improved tolerance of oxidative DNA damage

The pronounced peripheral vasoconstriction evoked by the diving response restricts blood supply to peripheral tissues of the diving mammal, causing an oxygen shortage (ischemia). Restoration of blood flow (reperfusion) to these tissues causes the production of reactive oxygen species (ROS) (*62*), which can damage DNA. Diving mammals are better adapted to tolerate frequent ischemia/reperfusion-induced ROS generation by possessing high levels of antioxidants (*63*). In addition to these increased antioxidant levels, we detected the inactivation of *POLM* (DNA polymerase mu) in the cetacean stem lineage (Figures 2A, S4), which has implications for improved tolerance of oxidative DNA lesions.

**Figure 2:**
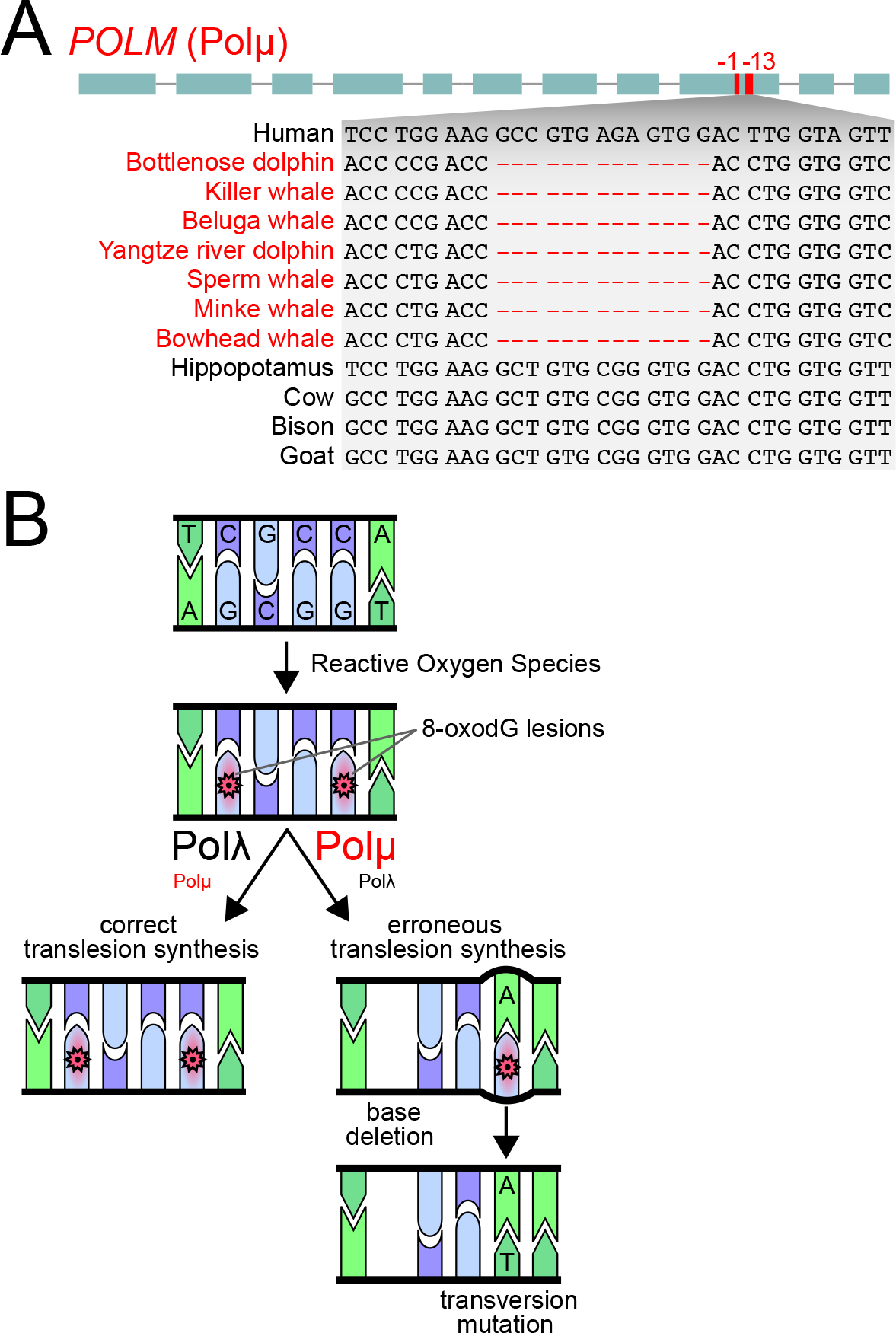
Loss of an error-prone DNA repair polymerase could have improved tolerance of oxidative DNA damage in cetaceans. (A) *POLM* was lost in the cetacean stem lineage, as shown by shared gene-inactivating mutations. Visualization as in Figure 1A. All inactivating mutations are shown in Figure S4. (B) Reactive Oxygen Species induce DNA damage, which includes oxidation of guanine (8-oxodG) as one the most frequent lesions. *POLM* encodes the DNA repair polymerase Polμ, which often does not perform a correct translesion synthesis (left) but instead introduces errors (right). In particular, Polμ typically deletes bases (*68*) or erroneously incorporates deoxy-adenosine opposite to 8-oxodG (instead of the correct deoxy-cytosine), which results in a C:G to A:T transversion mutation (*67*). In contrast to Polμ, another DNA repair polymerase Polλ is much less error-prone (*69*). Loss of *POLM* in cetaceans may have reduced the mutagenic potential of diving-induced oxidative stress by increasing the utilization of the more precise Polλ and accurate homology-directed DNA repair.

*POLM* encodes the DNA polymerase Polµ, which plays an integral role in DNA damage repair (*64*). The most severe type of DNA damage caused by ROS is a DNA double strand break (*65*). One mechanism to repair double strand breaks is non-homologous end joining (NHEJ), a process that ligates DNA strands without requiring a homologous template and re-synthesizes missing DNA bases by DNA polymerases. Polμ is able to direct synthesis across a variety of broken DNA backbone types, including ends that lack any complementarity (*66*). Importantly, this high flexibility comes at the cost of making Polµ more error-prone than Polλ, the second DNA polymerase that participates in NHEJ (*66*). One of the most frequent types of DNA damage caused by ROS is the oxidation of guanine, creating 8-oxodG (*67*). 8-oxodG is highly mutagenic as the bypassing Polµ resolves this lesion either by deleting bases (*68*) or by creating a transversion mutation (*67*) (Figure 2B). In contrast to Polµ, Polλ performs translesion synthesis with a much lower error rate (*69*).

Hence, the error-prone DNA repair polymerase Polµ likely constitutes a mutagenic risk factor in a regime of frequent oxidative stress, as experienced by diving cetaceans. Inactivation of Polµ in the cetacean stem lineage may have enhanced the fidelity of bypassing 8-oxodG lesions and repairing double-stranded breaks by increased utilization of the more precise Polλ, which is supported by mouse experiments. Compared with wild-type mice, *POLM* knockout mice showed significantly reduced mutagenic 8-oxodG translesion synthesis and exhibited a higher endurance when challenged with severe oxidative stress (*70*, *71*). *POLM* knockout mice also displayed improved learning abilities and greater liver regenerative capacity at high age, likely caused by delayed brain and liver aging due to a compensatory increase in accurate homology-directed DNA repair (*70*, *71*). Consequently, loss of *POLM* would have reduced the mutagenic potential of ROS, which readily form during repeated ischemia/reperfusion processes in diving mammals.

### Loss of lung-related genes and a high performance respiratory system

During diving, the cetacean lung collapses and reinflates during ascent. While lung collapse would represent a severe clinical problem for humans, it serves to reduce both buoyancy and the risk of developing decompression sickness in cetaceans (*72*). Our screen revealed the loss of two genes that have specific expression patterns in the lung, *MAP3K19* (mitogen-activated protein kinase 19) and *SEC14L3* (SEC14 like lipid binding 3) (Figures 3A,B, S5, S6).

**Figure 3:**
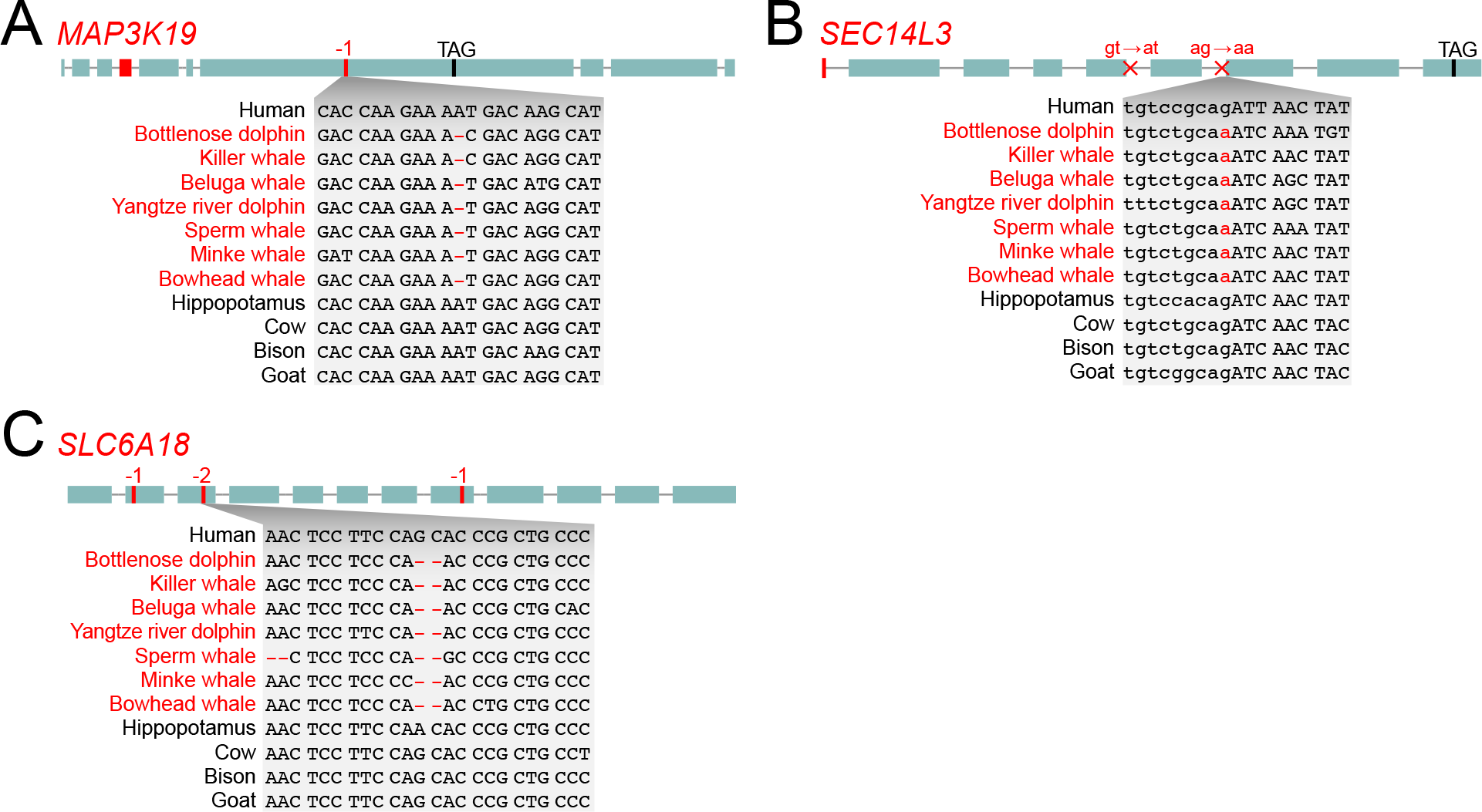
Loss of lung-related and renal transporter genes in the cetacean stem lineage. (A, B) The loss of *MAP3K19* and *SEC14L3*, which are specifically expressed in cell types of the lung, may relate to the high performance respiratory system of cetaceans. (C) The loss of the renal amino acid transporter *SLC6A18* offers an explanation for the low plasma arginine levels in cetaceans and may have contributed to stronger vasoconstriction during the diving response. Visualization as in Figure 1A. A shared donor (gt → at) and acceptor (ag → aa) splice site disrupting mutation is indicated in (B). All inactivating mutations are shown in Figures S5-S7.

*MAP3K19* is expressed in bronchial epithelial cells, type II pneumocytes and pulmonary macrophages (*73*). Overexpression of *MAP3K19* was detected in pulmonary macrophages of human patients suffering from idiopathic pulmonary fibrosis (*73*). This disease is believed to be caused by aberrant wound healing in response to injuries of the lung epithelium, leading to the abnormal accumulation of fibroblasts (fibrosis), excessive collagen secretion, and severely impaired lung function (*73*, *74*). Consistent with a fibrosis-promoting function of *MAP3K19*, inhibition of *MAP3K19* in mice protects from induced pulmonary fibrosis by significantly reducing fibrosis and collagen deposition (*73*). In a similar manner, *MAP3K19* loss may also have a protective effect in cetaceans where repeated lung collapse/reinflation events during deep dives causes shear forces that could increase the incidence of pulmonary microinjuries.

Furthermore, overexpression of *MAP3K19* was also detected in human patients suffering from chronic obstructive pulmonary disease (COPD) (*75*), a disease associated with cigarette smoking-induced oxidative stress. *MAP3K19* is upregulated in cells in response to oxidative and other types of environmental stress, and promotes the expression of pro-inflammatory chemokines (*75*). Further supporting a role of *MAP3K19* in the pathogenesis of COPD, inhibition of *MAP3K19* in mouse COPD models strongly reduced pulmonary inflammation and airway destruction (*75*). A hallmark of COPD is a reduction of alveolar elasticity caused by elastin degradation, which contributes to an incomplete emptying of the lung. Interestingly, cetaceans exhibit the opposite phenotype and have extensive elastic tissue in their lungs (*72*), which contributes to ‘explosive exhalation’, a breathing adaptation that allows renewal of ~90% of the air in the lung in a single breath (*76*). Therefore, similar to the previously-described loss of the elastin-degrading and COPD-overexpressed *MMP12* in aquatic mammals (*36*), the loss of *MAP3K19* may also be involved in the evolution of this breathing adaptation. More generally, the frequent oxidative stress faced by diving cetaceans, especially upon reoxygenation of the reinflated hypoxic lung, would increase the risk for *MAP3K19*-mediated chronic pulmonary inflammation and compromised respiratory function, which could have contributed to *MAP3K19* loss.

The second lung-expressed gene *SEC14L3* is expressed in airway ciliated cells and in alveolar type II cells that secrete pulmonary surfactant, the lipid-protein complex that prevents alveoli collapse (*77*, *78*). Similar to other surfactant-associated genes, *SEC14L3* expression is highly induced in the lungs before birth (*78*). SEC14L3 functions as a sensor of liposomal lipid-packing defects and may affect surfactant composition (*78*). In seals, the specific composition of pulmonary surfactant is thought to provide anti-adhesive properties, facilitating alveolar reinflation after collapse (*79*). Thus, it is possible that the loss of this surfactant-related gene in cetaceans is associated with frequent lung collapse and reinflation events in these diving mammals.

### Loss of a renal transporter gene and enhanced vasoconstriction during the diving response

Our screen revealed that cetaceans lost *SLC6A18* (Figures 3C, S7), which encodes a renal amino acid transporter that participates in reabsorption of arginine and other amino acids in the kidney proximal tubules. Knockout of *SLC6A18* in mice resulted in reduced plasma arginine levels (*80*). Thus, the loss of *SLC6A18* and its renal arginine reabsorbing activity provides one possible explanation for why cetaceans exhibit considerably lower plasma arginine levels in comparison to mice (*81*). In addition, *SLC6A18* knockout in mice resulted in stress-induced hypertension (*80*), a condition that involves vasoconstriction. This hypertension phenotype likely arises because lower arginine levels reduce the main substrate for the production of nitric oxide, a highly diffusible vasodilating substance (*80*). Consistently, *SLC6A18* inactivation caused persistent hypertension in a different mouse strain that is more susceptible to perturbations of nitric oxide production (*82*, *83*). This raises the possibility that the evolutionary loss of *SLC6A18* in the cetacean stem lineage may have contributed to an increased diving capacity by indirectly enhancing vasoconstriction during the diving response.

### Loss of an ion transporter gene and feeding in an aquatic environment

Saliva plays a role in lubricating the oral mucosa, in providing starch-degrading enzymes and in the perception of taste. All these functions became less important in an aquatic environment, where the abundance of water sufficiently lubricates food and dilutes salivary digestive enzymes. Additionally, the hyperosmotic marine environment necessitates strict housekeeping of freshwater resources in marine species (*84*), thus freshwater loss via saliva secretion may be detrimental. We found that *SLC4A9* (Solute Carrier Family 4 Member 9), a gene participating in saliva secretion, was lost in the cetacean stem lineage (Figures 4A, S8). Moreover, we found a convergent inactivation of this gene in the manatee, representing the only other fully-aquatic mammalian lineage (Figure S8).

**Figure 4:**
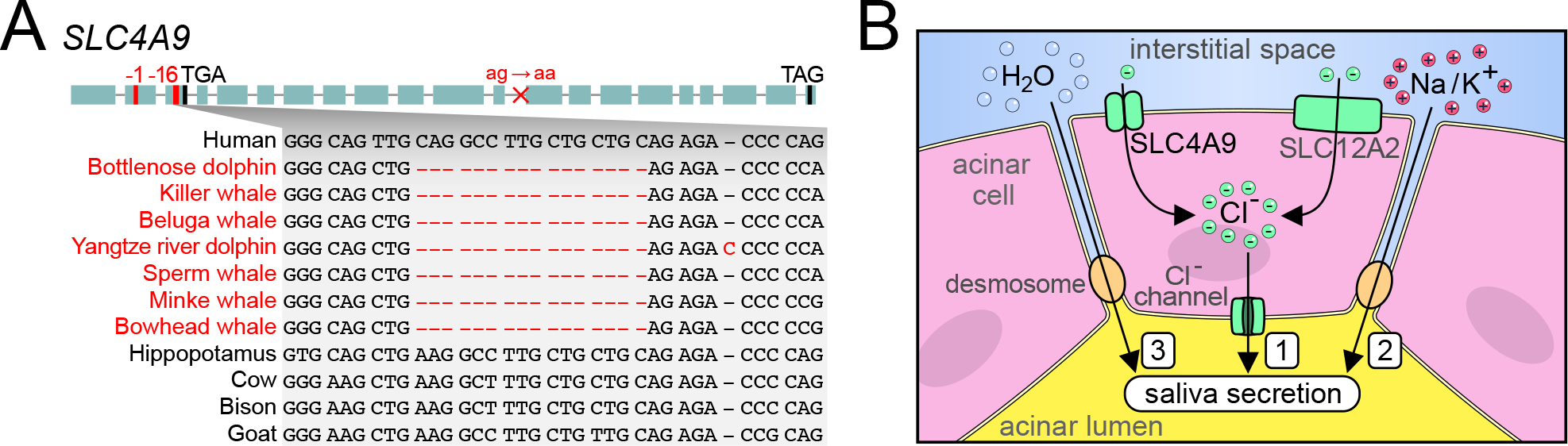
Loss of a pleiotropic ion transporter in cetaceans relates to the dispensability of saliva secretion. (A) Several shared inactivating mutations indicate that *SLC4A9* was lost in the cetacean stem lineage. Visualization as in Figure 1A. All inactivating mutations are shown in Figure S8. (B) Simplified illustration of saliva secretion. *SLC4A9* encodes an ion transporter. 1) In the submandibular salivary gland, SLC4A9 participates in creating a transepithelial chloride anion flux into the acinar lumen, together with another transporter SLC12A2 and chloride channels (*86*). 2) This first evokes a passive movement of cations across the tight junctions into the acinar lumen. 3) The resulting osmotic gradient induces a flow of water, which constitutes fluid secretion into the acinar lumen, the initial site of saliva secretion (*140*). *SLC4A9* knockout in mice leads to a 35% reduction in saliva secretion (*86*). The remaining saliva secretion potential in *SLC4A9* knockout mice is maintained by SLC12A2 (*86*). However, *SLC12A2* lacks inactivating mutations in cetaceans; such mutations lead to severe phenotypes in humans and mice (*141*, *142*), suggesting that gene essentiality maintained this gene in cetaceans. In addition to saliva secretion, SLC4A9 is also involved in transepithelial sodium ion flux in the kidney (not shown here) and participates in sodium chloride reabsorption (*90*), a process that is less important in hyperosmotic marine environments.

*SLC4A9* encodes an electroneutral ion exchange protein (*85*), which is expressed in the submandibular salivary gland. SLC4A9 is restricted to the basolateral membrane of acinar cells, where it participates in saliva secretion (*86*, *87*) (Figure 4B). *SLC4A9* knockout mice displayed a 35% reduction in saliva secreted from the submandibular gland (*86*). This suggests that loss of *SLC4A9* in cetaceans could contribute to a reduction of saliva secretion, which is in agreement with morphological observations that salivary glands are absent or atrophied in cetaceans (*88*, *89*).

In addition to the submandibular salivary gland, SLC4A9 is also expressed at the basolateral membrane of β-intercalated cells of the kidney, where it contributes to sodium chloride reabsorption (*90*). For species living in a hyperosmotic environment, where they incidentally ingest seawater with their prey, salt reabsorption by the kidney is probably less important (or even harmful) relative to efficient salt excretion. Thus, the loss of the salt reabsorbing factor *SLC4A9* may contribute to the high urinary concentrations of sodium and chloride in cetaceans as compared to cows (*91*). In summary, the pleiotropic *SLC4A9* gene was likely lost since both of its physiological processes – secretion of saliva and salt reabsorption – became dispensable in marine aquatic environments.

### Loss of melatonin biosynthesis/reception and the evolution unihemispheric sleep

Commitment to a fully-aquatic lifestyle also required distinct behavioral adaptations in stem cetaceans. Specifically, prolonged periods of sleep are obstructed by the needs to surface regularly to breathe and to constantly produce heat in the thermally-challenging environment of the ocean. Cetaceans are the only mammals thought to sleep exclusively unihemispherically, a type of sleep that allows one brain hemisphere to sleep while the awake hemisphere coordinates movement for surfacing and heat generation (*92*–*94*). Our screen uncovered that *AANAT* (aralkylamine N-acetyltransferase) was lost in the stem cetacean lineage (Figures 5A, S9). *AANAT* is a key gene required for synthesis of melatonin, the sleep hormone that influences wakefulness and circadian rhythms. Based on this observation, we inspected *ASMT* (acetylserotonin O-methyltransferase), encoding the second enzyme required for melatonin synthesis, and *MTNR1A* and *MTNR1B* (melatonin receptors 1A and 1B), encoding the two membrane-bound melatonin receptors. We found that all three genes were lost in all analyzed cetaceans, with *MTNR1B* being inactivated in the cetacean stem lineage (Figures 5A, S10), while *ASMT* and *MTNR1A* were probably inactivated independently after the split of odontocetes and mysticetes (Figures S11, S12). Thus, cetaceans have lost all genes required for melatonin biosynthesis and reception (Figure 5B). In line with these findings, cetaceans exhibit low levels of circulating melatonin, which does not follow a circadian pattern (*95*, *96*). Since dietary melatonin is readily transported into the blood stream (*97*), our finding that the melatonin-synthesizing enzymes *AANAT* and *ASMT* are lost in cetaceans further indicate that previously measured melatonin levels in cetaceans are not endogenous, but rather of dietary origin. Furthermore, the loss of the *ASMT* gene suggests that the previously-reported immunohistochemistry signal of ASMT protein in the retina, Harderian gland and gut of bottlenose dolphin (*96*) may be attributed to antibody cross-reactivity.

**Figure 5:**
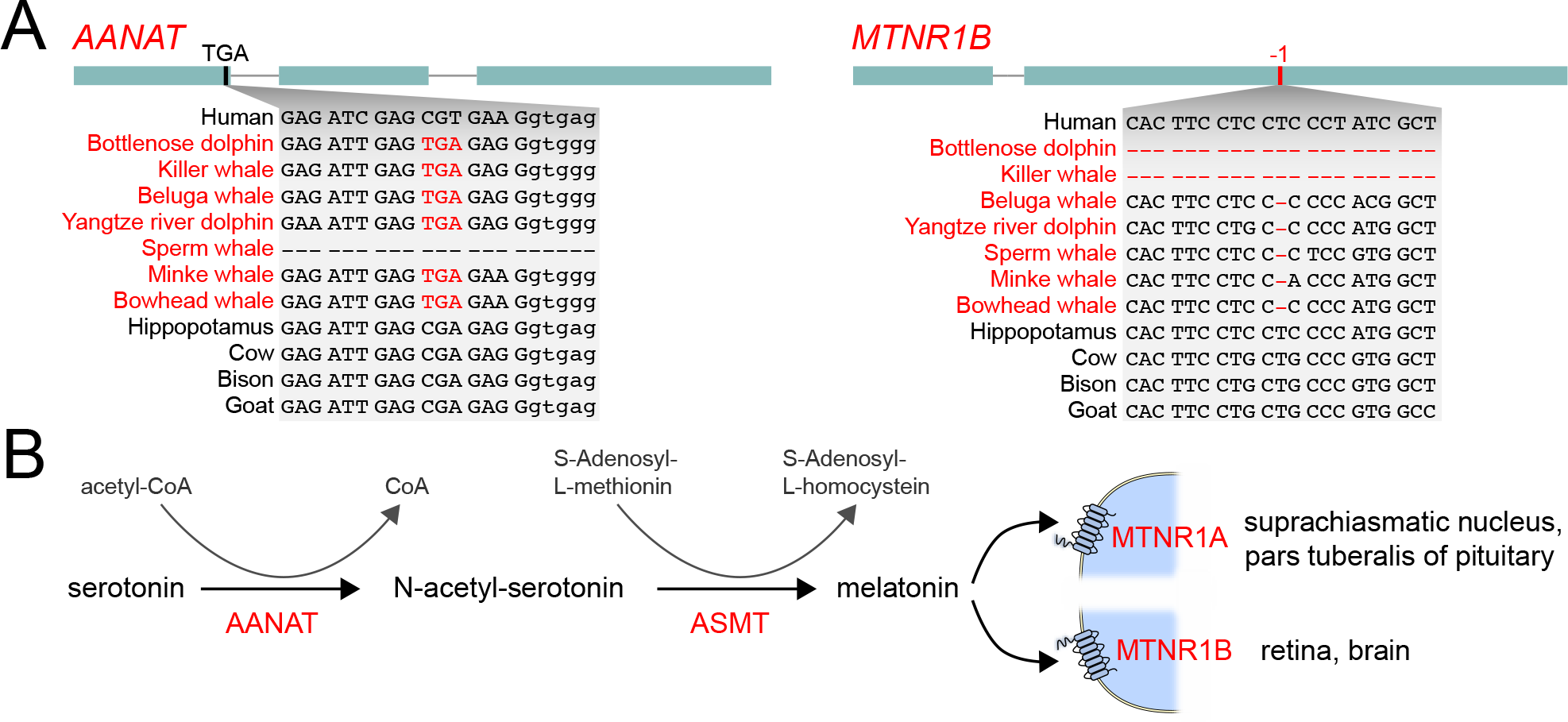
Complete loss of melatonin synthesis and reception may have been a precondition to exclusively adopt unihemispheric sleep in cetaceans. (A) Shared inactivating mutations indicate that *AANAT*, the first enzyme required to synthesize melatonin, and *MTNR1B*, one of the two melatonin receptors, were lost in the cetacean stem lineage. Subsequently, the second enzyme *ASMT* and the second receptor *MTNR1A* were probably independently lost in cetaceans after the split of odontocetes and mysticetes; however, overlapping deletions of the last *ASMT* coding exon and *MTNR1A* exon 2 do not exclude the possibility of ancestral gene losses. Visualization as in Figure 1A. All inactivating mutations are shown in Figures S9-S12. (B) Pathway to synthesize melatonin from serotonin and the main sites of expression of the two melatonin transmembrane receptors.

Melatonin is synthesized in the pineal gland in the absence of light (i.e. at night) by serial action of the enzymes AANAT and ASMT and thereby relays information on daytime and season (*98*, *99*). Polymorphisms in *AANAT* or *ASMT* impact sleep patterns in humans (*100*, *101*) and birds (*102*). Furthermore, knockout of *AANAT* in zebrafish decreased the length of sleep bouts, causing a ~50% reduction in nightly sleep time (*103*). It has been suggested that melatonin influences sleep-wake cycles mainly through binding the receptors encoded by *MTNR1A* and *MTNR1B* on cells of the suprachiasmatic nucleus (*104*, *105*). Accordingly, elimination of these two receptors significantly increased the time spent awake in mutant mice (*106*). Furthermore, a polymorphism in the promoter region of *MTNR1A* was linked to insomnia symptoms (*107*). In addition to influencing sleep, melatonin has also been shown to regulate core body temperature in a circadian manner (*108*), and high circulating melatonin levels evoke a reduction of core body temperature through increased distal heat loss (*109*).

Therefore, the potential benefits of abolishing melatonin production and reception for cetaceans were likely twofold. First, by helping to decouple sleep-wake patterns from daytime, loss of circadian melatonin production may have been a precondition to adopt unihemispheric sleep as the exclusive sleep pattern. Consistently, sleep in several cetacean species was observed to be equally distributed between day-and nighttime, and is thought to be primarily influenced by prey availability (*92*, *110*, *111*). Second, mechanisms that reduce core body temperature appear detrimental for species inhabiting a thermally challenging environment.

Interestingly, when we examined the melatonin biosynthesis/reception genes in the manatee, we found inactivating mutations in three of the four genes (*AANAT*, *ASMT* and *MTNR1B*, Figures S9-S11). Similar to unihemispheric sleep, manatees also display considerable interhemispheric asymmetry during slow wave sleep (*112*). Additionally, manatees seem to lack a pineal gland (*113*).

The results of our genomic analysis also have implications for the conflicting results on the morphological presence of the pineal gland in cetaceans. This gland has been reported to be absent or rudimentary in several cetaceans (but its absence can sometimes be variable between different individuals), while other species such as beluga, harbour porpoise and sperm whale appear to have a fully-developed pineal gland (*96*, *114*, *115*). Even if a pineal gland is present in some cetacean individuals or species, inactivating mutations in melatonin synthesis and receptor genes preclude a role for this gland in melatonin-mediated circadian rhythms.

### Loss of genes involved in immune system, muscle function, metabolism and development

Despite the fact that cetacean phenotypes have been extensively studied, our genomic screen for genes lost in the cetacean stem lineage detected several gene losses that imply changes in particular phenotypes, which have not been well characterized. For example, we found losses of genes involved in defense to infectious agents such as bacteria and viruses (*TRIM14*, *TREM1*; Figures S13, S14, Table S1). Furthermore, while mammals generally possess four genes encoding peptidoglycan recognition proteins, which are receptors important for antimicrobial function and for maintaining a healthy gut microbiome (*116*, *117*), cetaceans have lost three of these four genes (*PGLYRP1/3/4*, Figures S15-S17). While the loss of these genes highlights differences in the cetacean immune system, it is not clear whether these losses are potentially related to different pathogens encountered in a fully-aquatic environment, changes in gut microbiome composition in these obligate carnivores, or other reasons. Another example is *MSS51*, a gene that is predominantly expressed in fast glycolytic fibers of skeletal muscle (*118*). Inactivation of *MSS51* in muscle cell lines directs muscle energy metabolism towards beta-oxidation of fatty acids (*119*). *MSS51* was lost in the cetacean stem lineage (Figure S18), suggesting that muscle metabolism may be largely fueled by fatty acids, which would be consistent with a high intramuscular lipid content in cetaceans (*120*). Cetaceans also lost *ACSM3* (Figure S19), a gene involved in oxidation of the short chain fatty acid butyrate (*121*), but it is not clear if this loss relates to their carbohydrate-poor diet. The loss of *ADH4* (Figure S20) suggests differences in vitamin A metabolism (*122*). Finally, the cetacean loss of *SPINK7* (Figure S21), a gene involved in esophageal epithelium development, could be linked to the specific ontogeny of the cetacean esophagus, which is homologous to a ruminant’s forestomach (*123*). Interestingly, several of these genes are also convergently inactivated in the manatee (Table S1). Overall, this highlights the need for further studies to investigate how the loss of these genes affects immunity, metabolism and development in cetaceans.

### Loss of less well-characterized genes in cetaceans

Finally, we detected losses of genes that have no experimentally characterized function (Table S1). Some of these genes have tissue-specific expression patterns, exemplified by *FABP12* (Figure S22), a member of the fatty acid-binding protein family that is expressed in retina and testis of rats (*124*), *ASIC5* (Figure S23), an orphan acid-sensing ion channel specifically expressed in interneuron subtypes of the vestibulocerebellum that regulates balance and eye movement (*125*), or *C10orf82* (Figure S24), which is specifically expressed in the human testis (*126*). Natural losses of such uncharacterized genes provide intriguing candidates for future functional studies, which may help to relate evolutionary gene losses to particular cetacean phenotypes.

### Summary

By conducting a systematic screen for coding genes that were inactivated in the cetacean stem lineage, we discovered 85 gene losses, most of which have not been reported before. Many of these gene losses were likely neutral and their loss happened because of relaxed selection to maintain their function. Interestingly, this ‘use it or lose it’ principle may also apply to pleiotropic genes that are involved in more than one process. An example is the loss of the pleiotropic *SLC4A9* gene, which was likely permitted in cetaceans because both of its functions (saliva secretion, renal salt reabsorption) became less important in marine environments. Together with *KLK8*, a pleiotropic gene with epidermal and hippocampal functions that is convergently lost in cetaceans and manatees (*31*), this adds to a rather small list of known pleiotropic gene losses (*127*).

In addition to likely neutral gene losses, some of the genes that were lost in the cetacean stem lineage could have contributed to adapting to an aquatic environment, in particular in relation to the challenges of diving. The loss of *F12* and *KLKB1* likely reduced the risk of thrombus formation during diving. The loss of *POLM* likely reduced the mutagenic potential of ROS by indirectly enhancing the fidelity of oxidative DNA damage repair. The loss of *MAP3K19* protects from pulmonary fibrosis and from lung inflammation induced by oxidative stress. Since ischemia followed by reperfusion during diving generates ROS, losing these two genes may have contributed to better tolerating frequent diving-induced oxidative stress. *SLC6A18* loss could be involved in reduced plasma arginine levels, and thus indirectly enhance the vasoconstriction capacity during diving by reducing the substrate for synthesis of the potent vasodilator nitric oxide. Finally, the composition of pulmonary surfactant is important to allow lung reinflation after deep diving-induced alveolar collapse, which makes it interesting to investigate whether the loss of *SEC14L3* affects surfactant composition in cetaceans.

In conclusion, our findings suggest that gene losses in cetaceans are not only associated with aquatic specializations, but could have been involved in adapting to a fully-aquatic environment, which further supports that loss of ancestral genes can be a mechanism for phenotypic adaptation (*35*, *36*, *128*). More generally, our study highlights important genomic changes that occurred during the transition from land to water in cetaceans, and thus helps to understand the molecular determinants of their remarkable adaptations.

## Materials and Methods

### Detecting genes lost in cetaceans during the transition from land to water

We first searched for genes that exhibit inactivating mutations in the bottlenose dolphin, killer whale, sperm whale and common minke whale, using data generated by a previously-developed gene loss detection approach (*36*). Briefly, this approach used the human Ensembl version 90 gene annotation (*129*) and a genome alignment with the human hg38 assembly as the reference (*130*) (all analyzed assemblies are listed in Table S2) to detect stop codon mutations, frameshifting insertions or deletions, deletions of entire exons or genes, and mutations that disrupt splice sites (*36*). The approach performs a series of filter steps to remove artifacts related to genome assembly or alignment and evolutionary exon-intron structure changes in conserved genes. These steps comprise excluding those deletions that overlap assembly gaps in a query genome (*131*), re-aligning all coding exons with CESAR to detect evolutionary splice site shifts and to avoid spurious frameshifts due to alignment ambiguities (*132*, *133*), and excluding alignments to paralogs or processed pseudogenes. Finally, the approach considers all principal or alternative APPRIS isoforms of a gene (*134*) and outputs data for the isoform with the smallest number of inactivating mutations.

To screen for genes that were inactivated before the split of odontocetes and mysticetes, we extracted those genes with stop codon, frameshift or splice site mutations that are shared between species from both clades. We classified genes that are partially or completely deleted in odontocetes and mysticetes into two groups, those that have shared deletion breakpoints between at least one toothed and baleen whale and those where the deletion breakpoints are not shared between both lineages. To assess the deletion breakpoints up-and downstream of the deleted genes, we manually inspected the pairwise genome alignment chains (*135*) between human and cetaceans in the UCSC genome browser (*136*). We only included genes that exhibit shared breakpoints (such as *KLKB1*) since the most parsimonious explanation is a single deletion event in the cetacean stem lineage. However, it should be noted that a single ancestral deletion event may have been obscured by subsequent decay of the breakpoint regions in individual lineages, thus gene deletions without shared breakpoints, which are not included in this study, may have also occurred in the cetacean stem lineage. Likewise, we also excluded genes that exhibit smaller (stop codon, frameshift, splice site) inactivating mutations in one clade and are deleted in the other clade, even though the deletion may have happened after a single gene inactivation event occurred in the cetacean stem lineage. Since our alignment chains are sensitive enough to capture many gene duplications that happened before the divergence of mammals, we further used the alignment chains to exclude those gene loss candidates that likely possess another intact copy elsewhere in a cetacean genome due to a more recent gene duplication event. Finally, we excluded all genes that were also classified as lost in more than three terrestrial mammal species, as these genes are likely not involved in the adoption of a fully-aquatic lifestyle.

### Excluding genes with inactivating mutations in the hippopotamus lineage

To detect those genes that were inactivated during the transition from land to water in the cetacean stem lineage, we next excluded all genes that have inactivating mutations in the hippopotamus lineage. To this end, we first aligned the genome of the common hippopotamus (*40*) to the human hg38 assembly using lastz (parameters K = 2400, L = 3000 and default scoring matrix), axtChain, chainCleaner, and chainNet (all with default parameters) (*135*, *137*, *138*), and used our approach to detect inactivating mutations in the common hippopotamus.

For the eleven genes discussed in detail above (*F12*, *KLKB1*, *POLM*, *MAP3K19*, *SEC14L3*, *SLC6A18*, *SLC4A9*, *AANAT*, *ASMT*, *MTNR1A*, *MTNR1B*), we further corroborated the lack of inactivating mutations in the hippopotamus lineage. To this end, we made use of unassembled sequencing reads of the pygmy hippopotamus (*Choeropsis liberiensis*). We mapped these reads to exonic sequences of the common hippopotamus, including ~60 bp of flanking intron on each side, using Geneious 11.1.5 (*139*). In cases where an exon was not present in the common hippopotamus due to an assembly gap, we used the orthologous exon from cow and BLASTed the Sequence Read Archive (SRA) of the common hippopotamus (SRR5663647) to recover and assemble the missing exons. Manual inspection confirmed that all of the intact exons in the common hippopotamus also lack inactivating mutations in the pygmy hippopotamus.

### Validating gene loss in additional cetacean species

For all genes listed in Table S1, we investigated whether shared inactivating mutations are present in the genomes of three additional cetaceans that were not part of the whole-genome alignment (*130*), the Yangtze river dolphin, beluga whale and bowhead whale (*23*, *42*, *43*). To this end, we aligned these genomes to the hg38 assembly as described above and manually confirmed the presence of shared inactivation mutations.

## Data availability

The data supporting the findings of this study are publicly available or available within the Supplementary Information files. Specifically, all analyzed genome assemblies (Table S2) are publicly available on the UCSC genome browser and from NCBI. The list of 85 genes lost in the cetacean stem lineage together with functional annotations are provided in Table S1.

## Competing interests

The authors have no competing interests.

## Acknowledgment

We thank the genomics community for sequencing and assembling the genomes and the UCSC genome browser group for providing software and genome annotations. We also thank J. G. Roscito for help with gene annotations, G. Amato (NYZS) for providing a tissue sample for pygmy hippo genome sequencing, M. Collin for help with laboratory work, and the Computer Service Facilities of the MPI-CBG and MPI-PKS for their support. This work was supported by the Max Planck Society, the German Research Foundation (HI1423/3-1), the Leibniz Association (SAW-2016-SGN-2), and National Science Foundation (USA) grant (DEB-1457735).

